# Mice infected with *Mycobacterium tuberculosis* are resistant to secondary infection with SARS-CoV-2

**DOI:** 10.1101/2021.11.09.467862

**Authors:** Oscar Rosas Mejia, Erin S. Gloag, Jianying Li, Marisa Ruane-Foster, Tiffany A. Claeys, Daniela Farkas, Laszlo Farkas, Gang Xin, Richard T. Robinson

## Abstract

*Mycobacterium tuberculosis* (Mtb) and SARS-CoV-2 (CoV2) are the leading causes of death due to infectious disease. Although Mtb and CoV2 both cause serious and sometimes fatal respiratory infections, the effect of Mtb infection and its associated immune response on secondary infection with CoV2 is unknown. To address this question we applied two mouse models of COVID19, using mice which were chronically infected with Mtb. In both model systems, Mtb-infected mice were resistant to secondary CoV2 infection and its pathological consequences, and CoV2 infection did not affect Mtb burdens. Single cell RNA sequencing of coinfected and monoinfected lungs demonstrated the resistance of Mtb-infected mice is associated with expansion of T and B cell subsets upon viral challenge. Collectively, these data demonstrate that Mtb infection conditions the lung environment in a manner that is not conducive to CoV2 survival.

**AUTHOR SUMMARY:** *Mycobacterium tuberculosis* (Mtb) and SARS-CoV-2 (CoV2) are distinct organisms which both cause lung disease. We report the surprising observation that Mtb-infected mice are resistant to secondary infection with CoV2, with no impact on Mtb burden and resistance associating with lung T and B cell expansion.

## INTRODUCTION

The world is currently in the midst of two lung disease pandemics: COVID19 and tuberculosis (TB), the causative agents of which are SARS-CoV-2 (CoV2) and *Mycobacterium tuberculosis* (Mtb), respectively. Although COVID19 and TB both pose enormous health challenges, especially in countries where COVID19 vaccines are scarce, it unknown what if any effect Mtb infection has on host responses to CoV2 as there are few clinical reports of Mtb/CoV2 coinfection in the absence of other comorbidities [1, 2]. On the one hand, CoV2 infection may exacerbate the inflammatory response and pulmonary complications experienced by individuals with TB [3], analogous to that which is observed in the Mtb/Influenza A or Mtb/CMV coinfected individuals [4–7]. On the other hand, there is an inverse relationship between TB incidence rates and COVID19 mortality in numerous countries [8], and *Mycobacterium* spp express several proteins homologous to CoV2 antigens [9–11], raising the possibility that adaptive immune responses to Mtb may confer heterologous immunity against CoV2. To definitively address whether Mtb-infection impacts CoV2 elicited lung disease in a controlled setting, we applied two mouse models of COVID19 (CoV2 infection of K18-hACE2 mice [12], and mouse-adapted CoV2 [MACoV2] infection of C57BL/6 mice [13]), using mice that were chronically infected with Mtb. The results below support a model wherein Mtb infection confers resistance to secondary infection with CoV2 and its pathological consequences. The implications of these data for our understanding of COVID19 susceptibility and the limitations of our study are discussed.

## RESULTS

Details regarding the origin, culture, preparation and authentication of CoV2 (strain USA-WA1/2020), MACoV2 (strain MA10) and Mtb (strain H37Rv) are provided in our *Methods*. To determine if host responses to CoV2 are affected by Mtb-infection, K18-hACE2 (ACE2) and C57BL/6 (B6) mice were infected with low dose Mtb (~90 CFU) via aerosol delivery; thirty days later, the ACE2 mice were challenged with CoV2 (~25K PFU) via intranasal delivery (**FIG 1A**). These Mtb/CoV2 co-infected (Mtb^POS^CoV2^POS^) ACE2 mice were monitored daily for changes in weight, as were two control groups: ACE2 mice which were Mtb-infected at the same time (Day −30) but challenged with sterile media (Mtb^POS^CoV2^NEG^), and ACE2 mice which were not Mtb-infected prior to CoV2 challenge (Mtb^NEG^CoV2^POS^). On post-challenge Days 4, 7 and 14, groups of mice were euthanized and the lungs and other tissues were removed to assess Mtb and CoV2 burdens, as well as a number of immunological readouts. All mice were identically housed for the duration of the entire experiment. As anticipated, Mtb^NEG^CoV2^POS^ ACE2 mice lost a significant portion of body weight by post-challenge Day 7 (≤20%) (**FIG 1B**). Mtb^POS^CoV2^POS^ ACE2 mice, however, did not lose significant body weight and were otherwise indistinguishable from Mtb^POS^CoV2^NEG^ controls (**FIG 1B**). On post-challenge Day 4, lung CoV2 burdens were lower in Mtb^POS^CoV2^POS^ mice relative to Mtb^NEG^CoV2^POS^ mice, as assessed by either plaque assay (**FIG 1C**) or CoV2 N protein measurement (**FIG 1D**). Challenge with CoV2 did not affect Mtb growth, as Mtb CFU burdens in Mtb^POS^CoV2^POS^ and Mtb^POS^CoV2^NEG^ lungs did not differ after challenge (**FIG 1E**), nor did they differ in spleen (**FIG 1F**) or liver (**FIG 1G**). Consistent with the above Mtb CFU results, the abundance of acid fast bacilli (AFB) was also similar between Mtb^POS^CoV2^POS^ and Mtb^POS^CoV2^NEG^ lungs (**FIG 1H**). Transgenic human ACE2 expression also does affect Mtb growth, as CFU burdens in Mtb^POS^CoV2^NEG^ ACE2 mice were indistinguishable from Mtb-infected B6 controls (**FIG 1E-G**).

**Figure 1.**
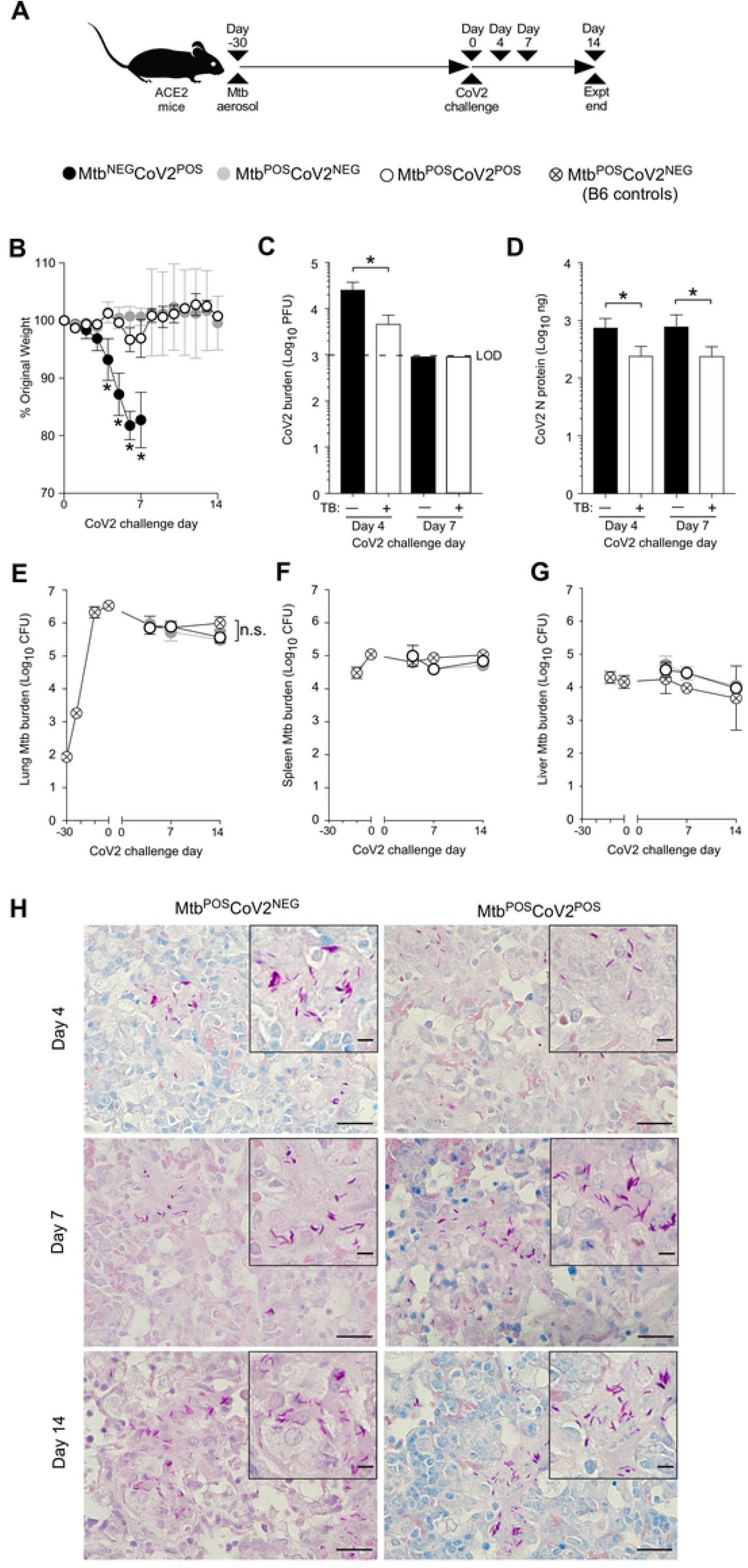
Mtb-infected ACE2 mice are resistant to secondary infection with CoV2. (**A**) Experimental overview of our ACE2:CoV2 model studies, wherein mice were infected via aerosol with Mtb (Day −30) and challenged 30 days later (Day 0) with CoV2. On post-challenge Day 4, Day 7 and Day 14, tissues were collected for microbiological and immunological assessments. Experimental groups included ACE2 mice which were not infected with Mtb prior to CoV2 challenge (Mtb^NEG^CoV2^POS^), ACE2 mice which were infected with Mtb but challenged with sterile saline (Mtb^POS^CoV2^NEG^), ACE2 mice which were infected with Mtb prior to CoV2 challenge (Mtb^POS^CoV2^POS^), and B6 controls which were infected with Mtb (to determine what if any impact human ACE2 transgene expression alone has on Mtb burdens). (**B**) The percent weight change experienced by each group of ACE2 mice following CoV2 challenge, as normalized to the original weight of each mouse. (**C**) CoV2 PFU burdens and (**D**) CoV2 N protein levels in the lungs of Mtb^NEG^CoV2^POS^ and Mtb^NEG^CoV2^POS^ mice. (**E-G**) Mtb CFU burdens in the (**E**) lungs, (**F**) spleen and (**G**) liver of Mtb^POS^CoV2^NEG^ and Mtb^POS^CoV2^POS^ mice, as well as B6 controls throughout the experiment time course. In each graph the following legend applies: Mtb^NEG^CoV2^POS^, black circles or bars; Mtb^POS^CoV2^NEG^, gray circles; Mtb^POS^CoV2^POS^, white circles or bars. (**H**) Representative micrographs of AFB stained lung sections, as collected from Mtb^POS^CoV2^NEG^ and Mtb^POS^CoV2^POS^ mice at the indicated times post-challenge. In each micrograph, the large scale bar is 20 μM and inset scale bar is 5 μm. This experiment was repeated twice, each with similar results (4 mice/group/timepoint). *, p ≤ 0.05 as determined by either Student’s t-test or ANOVA; n.s., not significant.

We next assessed the impact of Mtb infection on CoV2-elicited immune responses in the lung, using tissue from the same ACE2 transgenic mice described above. CoV2 infection elicits the expression of multiple inflammatory genes in mouse lungs [12]. Consistent with this, protein levels of IFNγ, IL6 and IL1β were elevated in Mtb^NEG^CoV2^POS^ lungs post-challenge Day 4 and/or Day 7, relative to uninfected (UI) controls (**FIG 2A-C**). In Mtb^POS^CoV2^NEG^ mice, lung protein levels of IFNγ, IL6 and IL1β were even higher, and were not affected by CoV2 challenge (**FIG2A-C**, compare Mtb^POS^CoV2^NEG^ and Mtb^POS^CoV2^POS^ levels). This pattern, wherein Mtb monoinfection induces high levels of a gene expression that are unchanged upon CoV2 challenge, was also observed for IFNγ and TNFα at the mRNA level (**FIG 2D-E**); however and notably, the resistance of Mtb^POS^CoV2^POS^ mice did not associate with elevated expression of the antiviral genes IFIT2 and IFIT3, which are otherwise induced in Mtb^NEG^CoV2^POS^ mice (**FIG 2F-G**), nor was CoV2 able to induce CCL2 expression in the presence of Mtb (**FIG 2H**). Expression of the anti-inflammatory cytokine IL10 was low in all experimental groups related to UI controls (**FIG 2I**).

**Figure 2.**
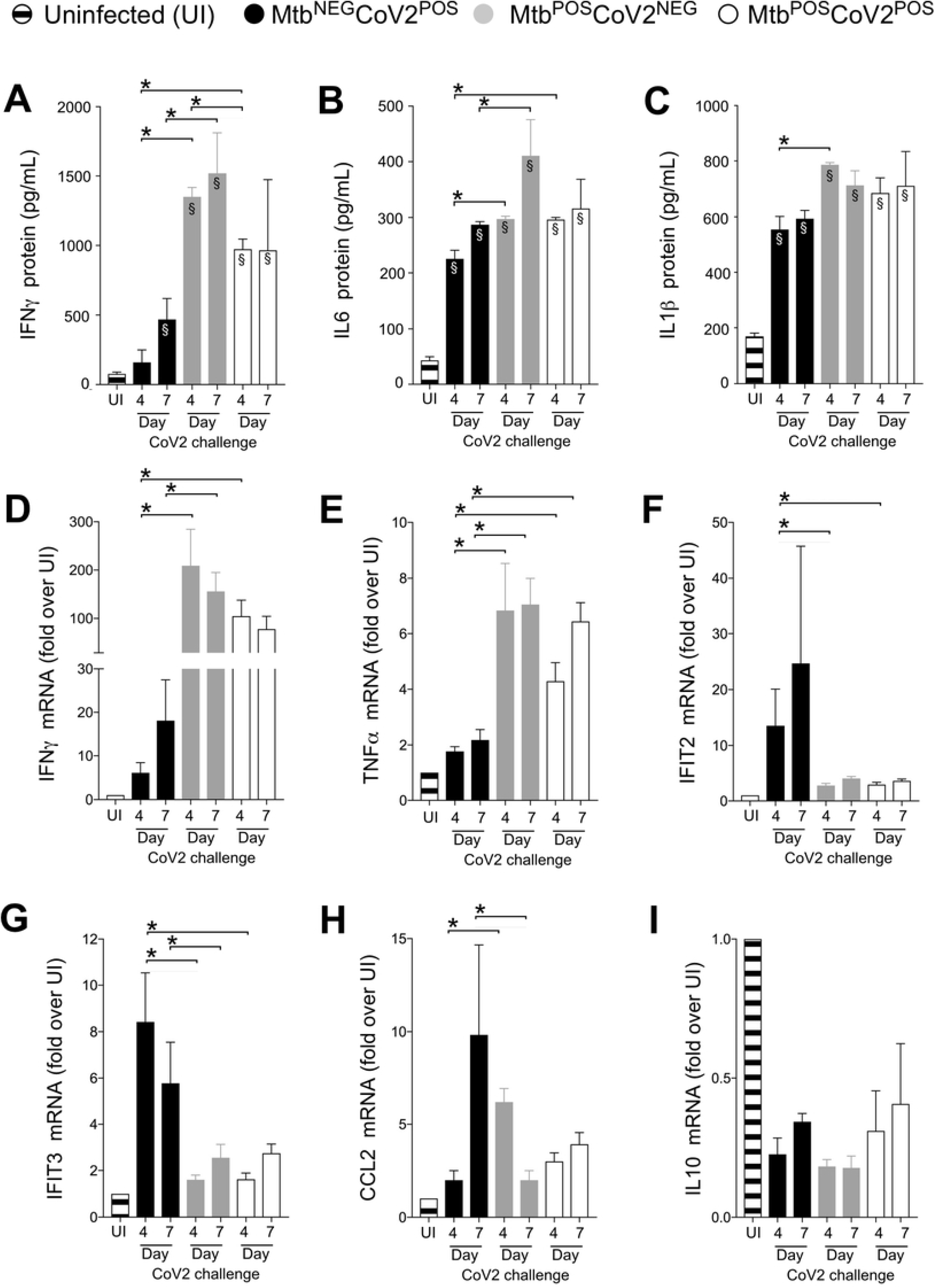
CoV2-elicted cytokine responses are muted in the presence of Mtb infection. On the indicated days, lung tissue from Mtb^NEG^CoV2^POS^, Mtb^POS^CoV2^NEG^, Mtb^POS^CoV2^POS^ and uninfected (UI) ACE2 mice was used to measure (**A-C**) protein levels of (**A**) IFNγ, (**B**) IL6 and (**C**) IL1β, as well as (**D-I**) mRNA levels of (**D**) IFNγ, (**E**) TNFα, (**F**) IFIT2, (**G**) IFIT3, (**H**) CCL2 and (**I**) IL10. This experiment was repeated twice, each with similar results (4 mice/group/timepoint). *, p ≤ 0.05 as determined by either Student’s t-test or ANOVA; §, significant relative to UI protein levels.

At a histological level, the lungs of Mtb^NEG^CoV2^POS^ mice exhibited a number of previously reported features [14] by post-challenge Day 4 (**FIG 3A**) and Day 7 (**FIG 3B**), including diffuse alveolar damage with inflammatory infiltrates and alveolar necrosis. Since these features were also observed in granulomatous lesions of Mtb^POS^CoV2^NEG^ lungs, a hallmark of Mtb infection, we could not use histology to observe whether Mtb inhibits CoV2-induced inflammation and alveolar necrosis. What could be observed, however, were differences between Mtb^NEG^CoV2^POS^ and Mtb^POS^CoV2^POS^ lungs with regards to hyaline membrane formation and pneumonia in the terminal bronchioles by Day 7 (**FIG 3B inset**), which were notable in Mtb^NEG^CoV2^POS^ lungs but absent from Mtb^POS^CoV2^POS^ lungs (pneumonia is not typical of Mtb-infected mice on the B6 background until ~1 year after infection [15]). Consistent with our assessment of lung CoV2 burdens (**FIG 1C-D**), anti-N protein immunohistochemistry (IHC) staining demonstrated fewer and less intense IHC+ regions within Mtb^POS^CoV2^POS^ lungs, relative to Mtb^NEG^CoV2^POS^ lungs (**FIG 3C-D**). Notably, the few IHC+ regions which were observed in Mtb^POS^CoV2^POS^ lungs were distal to granulomatous lesions that contain Mtb (**FIG 3C**).

**Figure 3.**
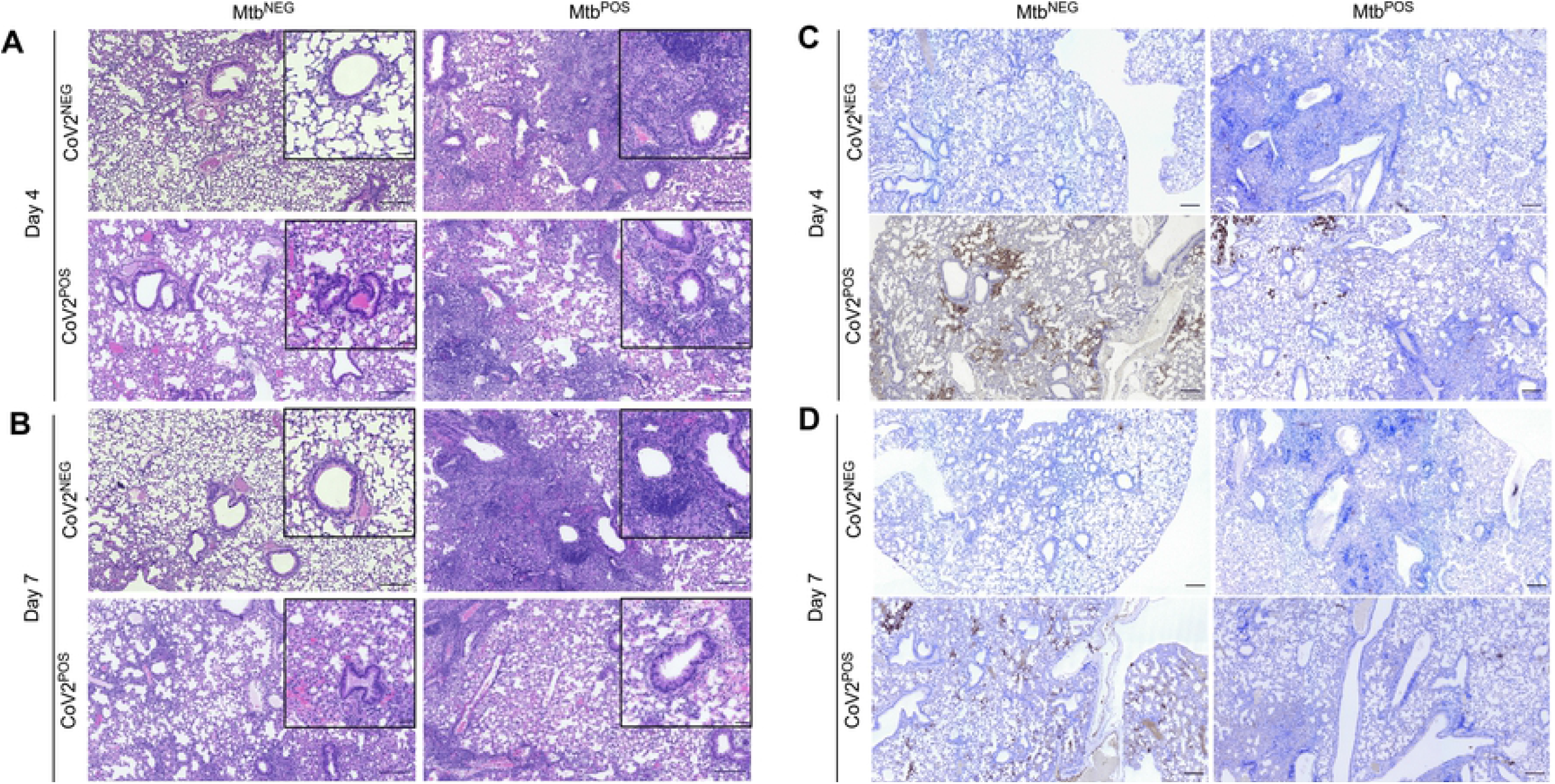
CoV2 infection of the airways and associated pneumonia are attenuated in the presence of Mtb. Representative micrographs of (**A-B**) H&E and (**C-D**) CoV2 N protein IHC stained lung sections from each experimental group, as collected on (**A, C**) Day 4 or (**B, D**) Day 7 post challenge. In each micrograph the large scale bar represents 200 microns; insets are 50 microns.

To determine whether Mtb-induced resistance to CoV2 was specific to the ACE2 transgenic model of COVID19, we performed the same set of experiments using a second mouse model of COVID19: MACoV2 infection of B6 mice [13]. As before, our experimental groups included B6 mice which were uninfected prior to MACoV2 challenge (Mtb^NEG^MACoV2^POS^), or Mtb-infected 30 days prior to challenge with MACoV2 (Mtb^POS^MACoV2^POS^) or vehicle control (Mtb^POS^MACoV2^NEG^) (**FIG 4A**). Whereas ACE2 mice which lost ≤20% body weight within 7 days of CoV2 challenge (**FIG 1B**), MACoV2 induced weight loss was less dramatic, with Mtb^NEG^MACoV2^POS^ mice losing ≤10% body weight within 7 days of MACoV2 challenge (**FIG 4B**). Nevertheless and consistent with our ACE2 model results, Mtb^POS^MACoV2^POS^ were resistant to MACoV2-elicited weight loss (**FIG 4B**), had lower viral burdens compared to Mtb^NEG^MACoV2^POS^ mice (**FIG 4C-D**) and no change in lung Mtb burdens following virus challenge (**FIG 4E**). Following virus challenge, Mtb^NEG^MACoV2^POS^ lungs exhibited transient increases in protein and mRNA levels of IFNγ (**FIG 4F-G**), IL6 (**FIG 4H**), IFIT3 (**FIG4I**), IFITM3 (**FIG4J**) and ACE2 (**FIG 4K**), consistent with previous reports of CoV2 inducing expression of its own receptor [16]. As was also observed in the ACE2 model (**FIG 2A-B**), protein levels of IFNγ and IL6 were already high in Mtb^POS^MACoV2^NEG^ lungs and unaffected by MACoV2 challenge (**FIG 4F, H**). MACoV2 elicited IFIT3 expression in both Mtb^NEG^MACoV2^POS^ and Mtb^POS^MACoV2^POS^ lungs, albeit lower in the latter group (**FIG 4J**). The resistance of Mtb-infected B6 mice to CoV2 was not attributable to an absence of lung ACE2 expression, as Mtb^POS^MACoV2^POS^ mice expressed higher than UI levels of ACE2 (**FIG 4K**); unlike Mtb^NEG^MACoV2^POS^ mice, however, ACE2 expression in Mtb^POS^MACoV2^POS^ lungs was not affected by MACoV2 challenge (**FIG 4K**).

**Figure 4.**
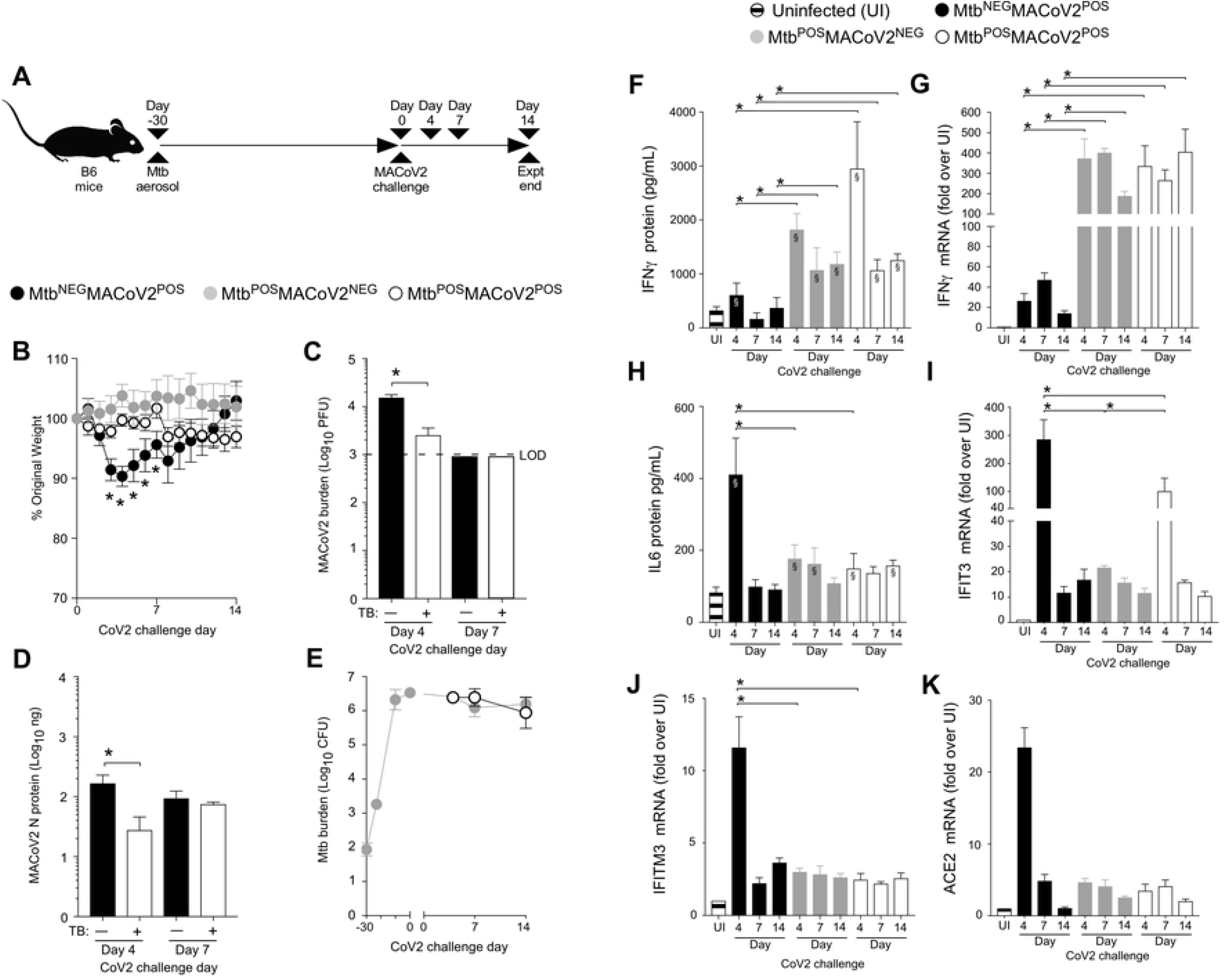
Mtb-infected B6 mice are resistant to secondary infection with MACoV2. (**A**) Experimental overview of our B6:MACoV2 model studies, wherein mice were infected via aerosol with Mtb (Day −30) and challenged 30 days later (Day 0) with MACoV2. On post-challenge Days 4, 7 and14 we collected lung tissue for microbiological and immunological assessments. Experimental groups included B6 mice which were not infected with Mtb prior to MACoV2 challenge (Mtb^NEG^MACoV2^POS^), B6 mice which were infected with Mtb but challenged with sterile saline (Mtb^POS^MACoV2^NEG^), and B6 mice which were infected with Mtb prior to CoV2 challenge (Mtb^POS^MACoV2^POS^). (**B**) The percent weight change experienced by each group of B6 mice following MACoV2 challenge, as normalized to the original weight of each mouse. (**C-D**) Lung viral burdens in Mtb^NEG^MACoV2^POS^ and Mtb^POS^MACoV2^POS^ mice, as measured by (**C**) MACoV2 PFU or (**D**) MACoV2 N protein concentration on the indicated days, as well as (**E**) lung Mtb CFU burdens at the same timepoints. (**F, H**) Lung protein levels of (**F**) IFNγ and (**H**) IL6, as well as (**G, I-K**) mRNA levels of (**G**) IFNγ, (**I**) IFIT3, (**J**) IFITM3 and (**K**) ACE2. This experiment was repeated twice, each with similar results (4 mice/group/timepoint). *, p ≤ 0.05 as determined by either Student’s t-test or ANOVA; §, significant relative to UI protein levels.

Finally, to discern the lung immune environment associated with MACoV2 resistance in Mtb-infected mice, we used scRNA-seq to analyze live CD45+ cells from the lungs of each group (UI, Mtb^NEG^MACoV2^POS^, Mtb^POS^MACoV2^NEG^ and Mtb^POS^MACoV2^POS^) on post-challenge Day 7. This timepoint enabled us to analyze immune cells after PFU are no longer detectable (**FIG 4C**). Live CD45+ cells were purified from the lungs of each group (4 mice per group) and used to prepare single-cell transcriptome datasets. These datasets separated into 12 clusters using the dimensionality reduction and clustering algorithms in the 10X Cell Ranger pipeline (**FIG 5A-C**). The expression profile of 20 myeloid and lymphoid lineage markers (*S100a4, S100a9, Cd8b1, Cd4, Cd79a, Ms4a1, Cybb, Mafb, Cd3g, Fcgr3, Cst3, Nme1, Itgam, Cd8a, Lig1, Ccna2, Ccr7, Il7r, Ncr1* and *Nkg7*) allowed us to assign biological identities to each cluster (**FIG 5D**). For each lineage marker, the average expression and percent positivity within each cluster were similar across all experimental groups (**SFIG1)**. We identified four T cell clusters (clusters 0, 2, 4 and 7), two B cell clusters (clusters 3 and 9), three myeloid-cell clusters (clusters 6, 8, and 10), one basophil cluster (cluster 11), one neutrophil cluster (cluster 5), and one natural killer–cell cluster (cluster 1) (**FIG 5A**). The extent to which these clusters were represented among all CD45+ cells varied by group (**FIG 5C, E**). We observed that innate clusters (i.e. NK, neutrophil, DC, MØ, CD11b+ MØ and basophils) comprised 50% of the UI lung, with T cells (42 %) and B cells (8%) making up the difference (**FIG 5D**). In Mtb^NEG^MACoV2^POS^ lungs, the representation of T cell (51 %) and B cell (16 %) clusters was higher, as were DC (4 %) and MØ (7 %) clusters. Relative to UI lungs, the Mtb^POS^MACoV2^NEG^ lung was characterized by the expansion of nearly all immune clusters (CD8 T cells, 8→15%; B cells, 6→13%; CD8 memory T cells, 7→9%; DCs, 2→4%; CD4 T cells, 3→6%; CD11b+ MØ, 1→2%; activated B cells, 2→4%; MØ, 1→5%) at the expense of neutrophils (17→7%), NK cells (28→15%) and naïve T cells (24→21%). Importantly, the profile of Mtb^POS^MACoV2^POS^ lungs closely resembled that of Mtb^POS^MACoV2^NEG^ lung, with the exception of expanded B cell (17%), CD8 memory T cell (10%), DC (6%) and activated B cell (5%) clusters, again at the expense of neutrophils, NK cells and naïve T cells (**FIG 5D**). Collectively, our scRNA seq data demonstrates the resistance of Mtb-infected mice to MACoV2 is associated with a lung immune environment that is largely similar to that observed in Mtb monoinfected lungs, with the exception of expanded T and B cell clusters.

**Figure 5.**
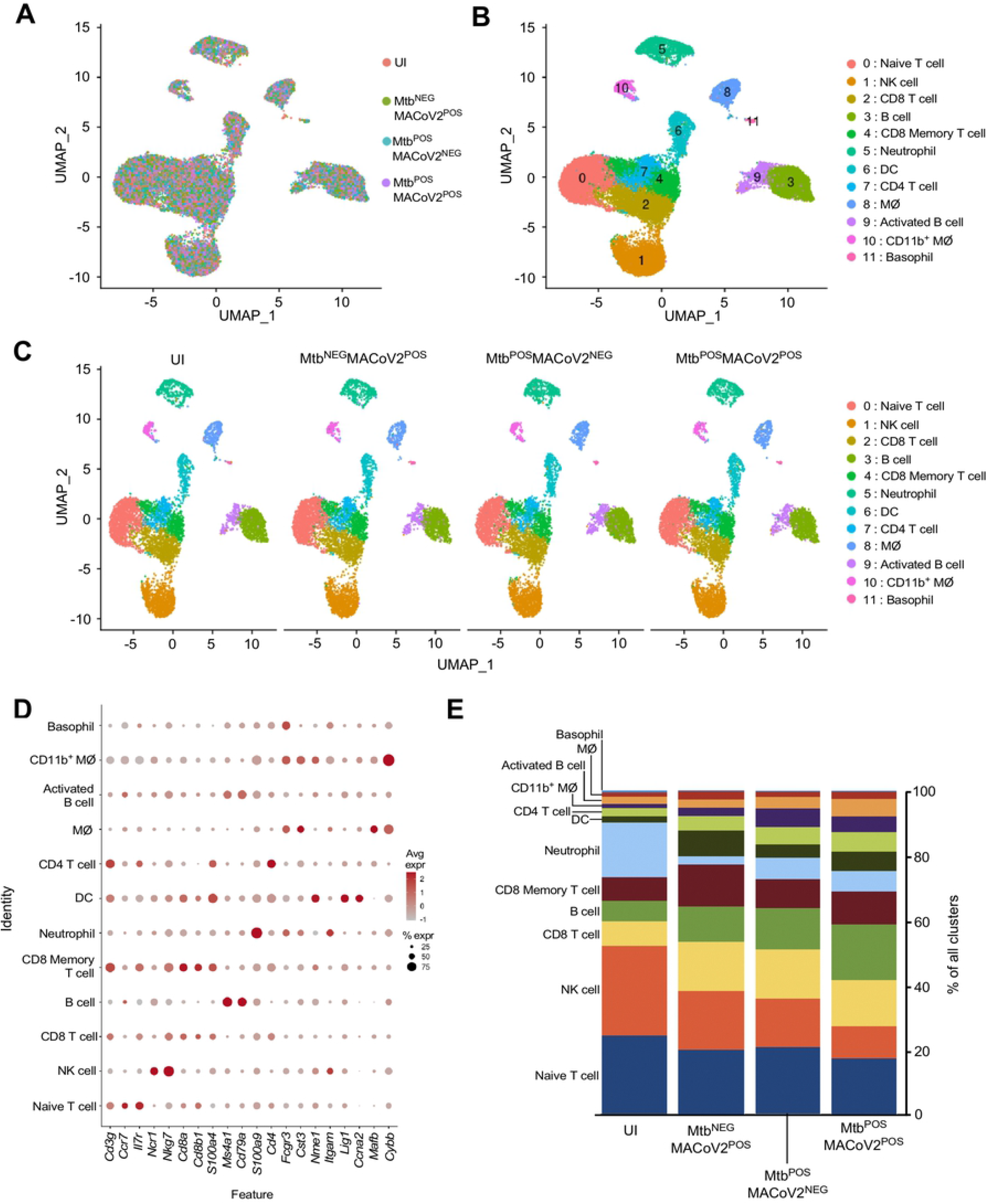
Lung T and B cell subsets expand upon challenge of Mtb^POS^ mice with MACoV2. As an unbiased means to define and compare the lung immune landscape, live CD45+ cells were purified from the lungs of four experimental groups (Uninfected, UI; Mtb^NEG^MACoV2^POS^; Mtb^POS^MACoV2^NEG^; Mtb^POS^MACoV2^POS^) on post-challenge Day 7 and used for scRNA analysis. (**A-C**) t-SNE plots of the resulting data, either (**A-B**) pooled across groups or (**C**) segregated by group to show (**A**) the extent of overlay and (**B-C**) clustering of data into 12 immune lineages. (**D**) The distribution and expression patterns of lineage defining genes which we used to annotate each cluster, as pooled from all group data (individual group data are shown in **Supplemental Figure 1**). (**E**) The proportion of each immune lineage in the lungs of each experimental group. MØ, macrophage; DC, dendritic cell; NK, natural killer.

## DISCUSSION

Our results demonstrate that Mtb infected mice are resistant to secondary infection with CoV2 and its pathological consequences. With regards to the mechanism of resistance, we believe the inflammatory nature of Mtb infection creates a lung environment that is inhospitable to CoV2 propagation. In the absence of Mtb, CoV2 enter cells via ACE2, propagates and triggers and inflammatory response that extends after CoV2 clearance and causes declines in lung function and death. In the presence of Mtb, CoV2 entry is likely unaffected since ACE2 is abundantly expressed in the Mtb-infected lung, but the extent of CoV2 propagation is low and the immunopathological responses typically triggered in mice (i.e. weight loss, pneumonia) are muted. This is likely due to one or both of the following reasons: (1) Mtb infected lungs already contain an array of immune innate lineages which restrict CoV2, or (2) Mtb elicits an adaptive immune response that cross reacts with CoV2 antigens and offers heterologous immunity. This latter explanation is supported by recent epidemiological studies of COVID among individuals vaccinated with *M. bovis* BCG [17–19], which depending on the strain has significant antigenic overlap with Mtb [20, 21]. The limitations of our study include its being performed in mice, which of course do not recapitulate all aspects of TB or COVID in humans, nor have we examined the long term impact of CoV2 on the host response to Mtb as we terminated our study fourteen days after CoV2 challenge. That said, we believe animal models of TB and COVID are ideal for studies of this nature because—if studies of COVID in individuals with other chronic lung diseases are any guide [22, 23]—it will likely be difficult to tease apart the impact of TB on COVID outcomes in humans given that individuals with TB often have numerous other comorbidities (e.g. malnourishment, HIV) that confound interpretation. Translated to human COVID susceptibility, our results suggest that individuals infected with Mtb generate an immune response that offers a degree of protection from subsequent or secondary infection with CoV2.

## MATERIALS & METHODS

### SARS-CoV-2 culture, preparation and authentication

All experiments involving SARS-CoV-2 followed procedures and protocols that are approved by The Ohio State University (OSU) Institutional Biosafety Committee. SARS-CoV-2, isolate USA-WA1/2020, was obtained from Biodefense and Emerging Infections Research Resources Repository (BEI Resources, Batch # 70034262). Mouse adapted SARS-CoV-2 variant strain MA10 [24] was likewise provided by BEI Resources (Cat # NR-55329). Virus was cultured, prepared and authenticated as we recently reported [25]. Namely, to establish the viral stocks used in our studies, a virus aliquot was thawed, diluted 1:10,000 in incomplete DMEM (Gibco; supplemented with 4.5 g/L D-glucose, 110 mg/L sodium pyruvate) and added to confluent VeroE6 cells (ATCC). Cells were incubated with virus for 1h (37°C, 5% CO_2_), after which time the media was replaced with complete DMEM (i.e. DMEM prepared as above, further supplemented with 4% heat-inactivated fetal bovine serum) and the cells were incubated for 3 days (37°C, 5% CO_2_) to allow virus propagation. After that period, visual inspection under light microscopy demonstrated near complete death of the infected VeroE6 cells. The supernatant was collected into 50mL conicals, centrifuged at low speed to remove cell debris and subsequently aliquoted, frozen and stored at −80°C. These frozen aliquots served as the stock tubes for all subsequent experiments. The live virus titer of our frozen aliquots was determined via the plaque assay described below. SARS-CoV-2 stocks were authenticated using a clinically validated clinical next-generation sequencing assay [26].

### *Mycobacterium tuberculosis* culture, preparation and authentication

All experiments involving *M. tuberculosis* (Mtb) followed procedures and protocols that are approved by The Ohio State University (OSU) Institutional Biosafety Committee. The virulent Mtb strain H37Rv (Trudeau Institute, Saranac Lake, NY) was grown in Proskauer Beck medium containing 0.05% tyloxapol to mid-log phase (37°C, 5% CO_2_) and frozen in 1-ml aliquots at −80°C. The live bacteria titer of our frozen aliquots was determined via plating serial dilutions on 7H11 agar media. To authenticate our Mtb stock we confirmed that the colony morphology, *in vitro* growth characteristics and *in vivo* virulence were consistent with our previous studies using the H37Rv strain [27].

### Mice

All mice were treated in accordance with OSU Institutional Animal Care and Use Committee (IACUC) guidelines and approved protocols. C57BL/6 and hemizygous K18-hACE C57BL/6J mice (strain: 2B6.Cg-Tg(K18-ACE2)2Prlmn/J) were purchased from Jackson Laboratory (Bar Harbor, ME) and housed at OSU within an AALAC-accredited facility (University Laboratory Animal Resources, ULAR).

### Aerosol Mtb infection

Mice were aerosol infected with Mtb H37Rv per our previous studies using the Glas-Col inhalation system [27]. For bacterial load determinations, the lungs, spleen, and liver were aseptically removed and individually homogenized in sterile normal saline (Gentle Macs system, program “RNA” run 2X). Serial dilutions of each organ were then plated on 7H11 and colonies counted after 2-3 weeks incubation at 37°C 5% CO_2_. Lungs from control mice were plated on post-infection Day 1 to verify the delivery of ~80 Mtb CFU.

### Intranasal CoV2 challenge

Mice that were either uninfected (UI) or previously infected with aerosol Mtb (Mtb^POS^) mice were challenged with either CoV2 or MACoV2. At the time of challenge, mice were anesthetized with isoflurane, weighed and held at a semi-supine position while 50 μL of CoV2-containing PBS (2.5 × 10^4^ PFU) or MACoV2 (2.5 × 10^4^ PFU) was given via intranasal (i.n.) instillation. Control mice were given the same volume of sterile PBS, using the same anesthesia and i.n. instillation protocol. After i.n. instillation, each mouse was returned to its home cage, house and monitored daily for changes in weight or body condition. For viral load determinations, the lungs of challenged animals were aseptically removed and individually homogenized as described above; serial dilutions were then used in the plaque assay described below.

### CoV2 plaque assay

A modified version of the plaque assay developed by the Diamond laboratory [28] was used to determine lung viral burdens in challenged animals, the details of which we have reported [29]. Namely, one day prior to the assay start we seeded 12-well with VeroE6 cells and incubated overnight (37°C 5% CO_2_) such that each well was confluent by the assay start. On the day of the assay, serial dilutions of virus-containing material (e.g. lung homogenate) were prepared in cDMEM and warmed to 37°C. Media from each well of the 12-well plate was gently removed via pipette and replaced with 500uL of each virus sample dilution, the volume pipetted down the side of the well so as not to disturb the VeroE6 monolayer. The plate was incubated for 1 hr at 37°C 5% CO_2_. During this incubation period, a solution comprising a 1:0.7 mixture of cDMEM and 2% methylcellulose (viscosity: 4000 cP) was freshly made and warmed to 37°C in a water bath. After the 1 hr incubation period was over, the supernatant was removed from each well and replaced with 1 mL of the pre-warmed cDMEM:methylcellulose mixture. The culture plate was then returned to the incubator and left undisturbed for 3 days. On the final day, the cDMEM:methylcellulose mixture was removed from each well, cells were fixed with 4% para-formaldehyde in PBS (20 minutes, room temperature), washed with PBS and stained with 0.05% crystal violet (in 20% methanol). After rinsing plates with distilled water, plates were dried, and plaques were counted under a light microscope.

### Histology

The inferior lung lobe was removed from mice and fixed in 10% formalin. Sample processing, paraffin embedding, H&E and acid fast bacilli (AFB) staining was performed by the OSU Comparative Pathology & Mouse Phenotyping Shared Resource (CPMPSR). Immunohistochemistry (IHC) was performed using a monoclonal antibody specific to SARS-CoV-2 Nucleocapsid (clone B46F; ThermoFisher) per previously reported methods [30]. Histology slides were imaged using a Nikon Ti2 widefield microscope fitted with 4x, 10x and 60x CFI Plan Fluor objectives and a DS-Fi3 color camera. Images were processed using FIJI [31] and compiled using BioRender.com

### ELISA

CoV2 N protein levels in lung homogenates were determined using a commercially available ELISA kit (ADS Biotec), as were protein levels of the cytokines IL1β, IL6 and IFNγ (Biolegend). ELISA kits were used per manufacturer protocols.

### Quantitative Real Time PCR

Lung RNA was extracted from the superior lung lobe using the RNeasy Mini Kit method (Qiagen) and reverse transcribed using the SuperScript VILO cDNA Synthesis Kit method (ThermoFisher). Quantitative real time PCR (qRT-PCR) was performed on a C1000 Touch Thermocycler (Bio-Rad) using SYBR Select Master Mix (Applied Biosystems) per manufacturer protocols. The primer sequences used to amplify cDNA for genes of interest were previously published [32, 33]. Each biological replicate was performed in technical duplicate and data were analyzed using the ΔΔCt method.

### Cell purification

To purify live CD45+ cells for single cell RNA sequencing, lungs from uninfected, Mtb- or MACoV2-monoinfected and Mtb/MACoV2 coinfected mice were removed and treated in an identical manner. Lungs were first digested in a DNase/collagenase mixture [34]; dead cells from the resulting slurry were then removed via negative magnetic selection using the Dead Cell Removal kit method (Miltenyi). The live cells were then mixed with CD45 microbeads (Miltenyi) and used for positive magnetic selection of live CD45+ cells. Trypan blue staining was used to confirm cell viability. Cells were the prepared for single cell partitioning via a 10X Genomics Chromium Controller using manufacturer provided protocols (10x Genomics Document Number CG000136). 1 x 10^4^ cells per experimental group were loaded onto the Controller and partitioned, as carried out by the OSU Genomics Shared Resource core.

### Single cell RNA sequencing (scRNA seq)

scRNA-seq libraries were prepared and analyzed using the 10X Genomics and Illumina platforms, respectively, per previously reported methods [35].

### Statistical analysis

All experiments were performed using randomly assigned mice without investigator blinding. All data points and *p* values reflect biological replicates from at least two independent experiments per figure (4 mice per group per timepoint). Statistical analysis was performed using GraphPad Prism. Unpaired, two-tailed Student t tests and one-way ANOVA tests with post hoc Tukey-Kramer corrections were used to assess statistical significance. Graphs were likewise generated in GraphPad Prism. The only exception to this were the t-distributed stochastic neighbor embedding (t-SNE), annotation and graphing associated with our scRNA analysis, which was performed with Cell Ranger and RStudio.

## ACKNOWLEDGEMENTS

The project described was supported by funds from the NIH/NIAID (R01AI121212 to RTR) and The Ohio State University (OSU). ORM was funded by an OSU Advancing Research in Infection and Immunity Fellowship Award; ESG was funded by an American Heart Association Career Development Award (19CDA34630005). We would like to acknowledge the laboratories of Jianrong Li (OSU), Mark Peeples (OSU & Nationwide Children’s Hospital) and Jacob Yount (OSU) who grew and provided the MACoV2 used in our studies, as well as BSL3 Director Luanne Hall-Stoodley and BSL3 Research Assistant Abigail Mayer for maintaining the facilities needed for our studies.

---

**SUPPLEMENTAL FIGURE 1. Lineage defining markers were similarly expressed across uninfected (UI), Mtb^NEG^MACoV2^POS^, Mtb^POS^MACoV2^NEG^ and Mtb^POS^MACoV2^POS^ groups.** The distribution and expression patterns of lineage defining genes that were used to annotate each t-SNE cluster, as shown for each individual experimental group (pooled group data are shown in **Figure 5**).

## REFERENCES

1. Riou C, du Bruyn E, Stek C, Daroowala R, Goliath RT, Abrahams F, et al. Relationship of SARS-CoV-2-specific CD4 response to COVID-19 severity and impact of HIV-1 and tuberculosis coinfection. J Clin Invest. 2021;131(12). Epub 2021/05/05. doi: 10.1172/JCI149125. PubMed PMID: 33945513; PubMed Central PMCID: PMCPMC8203446.

2. Tamuzi JL, Ayele BT, Shumba CS, Adetokunboh OO, Uwimana-Nicol J, Haile ZT, et al. Implications of COVID-19 in high burden countries for HIV/TB: A systematic review of evidence. BMC Infect Dis. 2020;20(1):744. Epub 2020/10/11. doi: 10.1186/s12879-020-05450-4. PubMed PMID: 33036570; PubMed Central PMCID: PMCPMC7545798.

3. Mousquer GT, Peres A, Fiegenbaum M. Pathology of TB/COVID-19 Co-Infection: The phantom menace. Tuberculosis (Edinb). 2021;126:102020. Epub 2020/11/28. doi: 10.1016/j.tube.2020.102020. PubMed PMID: 33246269; PubMed Central PMCID: PMCPMC7669479.

4. Mendy J, Jarju S, Heslop R, Bojang AL, Kampmann B, Sutherland JS. Changes in Mycobacterium tuberculosis-Specific Immunity With Influenza co-infection at Time of TB Diagnosis. Front Immunol. 2018;9:3093. Epub 2019/01/22. doi: 10.3389/fimmu.2018.03093. PubMed PMID: 30662443; PubMed Central PMCID: PMCPMC6328457.

5. Cobelens F, Nagelkerke N, Fletcher H. The convergent epidemiology of tuberculosis and human cytomegalovirus infection. F1000Res. 2018;7:280. Epub 2018/05/24. doi: 10.12688/f1000research.14184.2. PubMed PMID: 29780582; PubMed Central PMCID: PMCPMC5934687.

6. Walaza S, Tempia S, Dawood H, Variava E, Moyes J, Cohen AL, et al. Influenza virus infection is associated with increased risk of death amongst patients hospitalized with confirmed pulmonary tuberculosis in South Africa, 2010-2011. BMC Infect Dis. 2015;15:26. Epub 2015/01/28. doi: 10.1186/s12879-015-0746-x. PubMed PMID: 25623944; PubMed Central PMCID: PMCPMC4316613.

7. Redford PS, Mayer-Barber KD, McNab FW, Stavropoulos E, Wack A, Sher A, et al. Influenza A virus impairs control of Mycobacterium tuberculosis coinfection through a type I interferon receptor-dependent pathway. J Infect Dis. 2014;209(2):270–4. Epub 2013/08/13. doi: 10.1093/infdis/jit424. PubMed PMID: 23935205; PubMed Central PMCID: PMCPMC3873785.

8. Inoue K, Kashima S. Association of the past epidemic of Mycobacterium tuberculosis with mortality and incidence of COVID-19. PLoS One. 2021;16(6):e0253169. Epub 2021/06/19. doi: 10.1371/journal.pone.0253169. PubMed PMID: 34143810; PubMed Central PMCID: PMCPMC8213125.

9. Eggenhuizen PJ, Ng BH, Chang J, Fell AL, Cheong RMY, Wong WY, et al. BCG Vaccine Derived Peptides Induce SARS-CoV-2 T Cell Cross-Reactivity. Front Immunol. 2021;12:692729. Epub 2021/08/24. doi: 10.3389/fimmu.2021.692729. PubMed PMID: 34421902; PubMed Central PMCID: PMCPMC8374943.

10. Dakal TC. Antigenic sites in SARS-CoV-2 spike RBD show molecular similarity with pathogenic antigenic determinants and harbors peptides for vaccine development. Immunobiology. 2021;226(5):152091. Epub 2021/07/26. doi: 10.1016/j.imbio.2021.152091. PubMed PMID: 34303920; PubMed Central PMCID: PMCPMC8297981.

11. Nuovo G, Tili E, Suster D, Matys E, Hupp L, Magro C. Strong homology between SARS-CoV-2 envelope protein and a Mycobacterium sp. antigen allows rapid diagnosis of Mycobacterial infections and may provide specific anti-SARS-CoV-2 immunity via the BCG vaccine. Ann Diagn Pathol. 2020;48:151600. Epub 2020/08/18. doi: 10.1016/j.anndiagpath.2020.151600. PubMed PMID: 32805515; PubMed Central PMCID: PMCPMC7423587.

12. Winkler ES, Bailey AL, Kafai NM, Nair S, McCune BT, Yu J, et al. SARS-CoV-2 infection of human ACE2-transgenic mice causes severe lung inflammation and impaired function. Nat Immunol. 2020;21(11):1327–35. Epub 2020/08/26. doi: 10.1038/s41590-020-0778-2. PubMed PMID: 32839612; PubMed Central PMCID: PMCPMC7578095.

13. Dinnon KH, 3rd, Leist SR, Schafer A, Edwards CE, Martinez DR, Montgomery SA, et al. A mouse-adapted model of SARS-CoV-2 to test COVID-19 countermeasures. Nature. 2020;586(7830):560–6. Epub 2020/08/28. doi: 10.1038/s41586-020-2708-8. PubMed PMID: 32854108; PubMed Central PMCID: PMCPMC8034761.

14. Jiang RD, Liu MQ, Chen Y, Shan C, Zhou YW, Shen XR, et al. Pathogenesis of SARS-CoV-2 in Transgenic Mice Expressing Human Angiotensin-Converting Enzyme 2. Cell. 2020;182(1):50–8 e8. Epub 2020/06/10. doi: 10.1016/j.cell.2020.05.027. PubMed PMID: 32516571; PubMed Central PMCID: PMCPMC7241398.

15. Rhoades ER, Frank AA, Orme IM. Progression of chronic pulmonary tuberculosis in mice aerogenically infected with virulent Mycobacterium tuberculosis. Tuber Lung Dis. 1997;78(1):57–66. Epub 1997/01/01. doi: 10.1016/s0962-8479(97)90016-2. PubMed PMID: 9666963.

16. Ziegler CGK, Allon SJ, Nyquist SK, Mbano IM, Miao VN, Tzouanas CN, et al. SARS-CoV-2 Receptor ACE2 Is an Interferon-Stimulated Gene in Human Airway Epithelial Cells and Is Detected in Specific Cell Subsets across Tissues. Cell. 2020;181(5):1016–35 e19. Epub 2020/05/16. doi: 10.1016/j.cell.2020.04.035. PubMed PMID: 32413319; PubMed Central PMCID: PMCPMC7252096.

17. Escobar LE, Molina-Cruz A, Barillas-Mury C. BCG vaccine protection from severe coronavirus disease 2019 (COVID-19). Proc Natl Acad Sci U S A. 2020;117(30):17720–6. Epub 2020/07/11. doi: 10.1073/pnas.2008410117. PubMed PMID: 32647056; PubMed Central PMCID: PMCPMC7395502.

18. Hauer J, Fischer U, Auer F, Borkhardt A. Regional BCG vaccination policy in former East- and West Germany may impact on both severity of SARS-CoV-2 and incidence of childhood leukemia. Leukemia. 2020;34(8):2217–9. Epub 2020/06/20. doi: 10.1038/s41375-020-0871-4. PubMed PMID: 32555367; PubMed Central PMCID: PMCPMC7301049.

19. O’Neill LAJ, Netea MG. BCG-induced trained immunity: can it offer protection against COVID-19? Nat Rev Immunol. 2020;20(6):335–7. Epub 2020/05/13. doi: 10.1038/s41577-020-0337-y. PubMed PMID: 32393823; PubMed Central PMCID: PMCPMC7212510.

20. Coppola M, Jurion F, van den Eeden SJF, Tima HG, Franken K, Geluk A, et al. In-vivo expressed Mycobacterium tuberculosis antigens recognised in three mouse strains after infection and BCG vaccination. NPJ Vaccines. 2021;6(1):81. Epub 2021/06/05. doi: 10.1038/s41541-021-00343-2. PubMed PMID: 34083546; PubMed Central PMCID: PMCPMC8175414.

21. Zhang W, Zhang Y, Zheng H, Pan Y, Liu H, Du P, et al. Genome sequencing and analysis of BCG vaccine strains. PLoS One. 2013;8(8):e71243. Epub 2013/08/27. doi: 10.1371/journal.pone.0071243. PubMed PMID: 23977002; PubMed Central PMCID: PMCPMC3747166.

22. Lacedonia D, Scioscia G, Santomasi C, Fuso P, Carpagnano GE, Portacci A, et al. Impact of smoking, COPD and comorbidities on the mortality of COVID-19 patients. Sci Rep. 2021;11(1):19251. Epub 2021/09/30. doi: 10.1038/s41598-021-98749-4. PubMed PMID: 34584165; PubMed Central PMCID: PMCPMC8478875.

23. Hadi YB, Lakhani DA, Naqvi SFZ, Singh S, Kupec JT. Outcomes of SARS-CoV-2 infection in patients with pulmonary sarcoidosis: A multicenter retrospective research network study. Respir Med. 2021;187:106538. Epub 2021/07/30. doi: 10.1016/j.rmed.2021.106538. PubMed PMID: 34325226; PubMed Central PMCID: PMCPMC8297986.

24. Leist SR, Dinnon KH, 3rd, Schafer A, Tse LV, Okuda K, Hou YJ, et al. A Mouse-Adapted SARS-CoV-2 Induces Acute Lung Injury and Mortality in Standard Laboratory Mice. Cell. 2020;183(4):1070–85 e12. Epub 2020/10/09. doi: 10.1016/j.cell.2020.09.050. PubMed PMID: 33031744; PubMed Central PMCID: PMCPMC7510428.

25. Bednash JS, Kagan VE, Englert JA, Farkas D, Tyurina YY, Tyurin VA, et al. Syrian hamsters as a model of lung injury with SARS-CoV-2 infection: pathologic, physiologic and detailed molecular profiling. Transl Res. 2021. Epub 2021/11/07. doi: 10.1016/j.trsl.2021.10.007. PubMed PMID: 34740873.

26. Wang H, Jean S, Eltringham R, Madison J, Snyder P, Tu H, et al. Mutation-Specific SARS-CoV-2 PCR Screen: Rapid and Accurate Detection of Variants of Concern and the Identification of a Newly Emerging Variant with Spike L452R Mutation. J Clin Microbiol. 2021;59(8):e0092621. Epub 2021/05/21. doi: 10.1128/JCM.00926-21. PubMed PMID: 34011523; PubMed Central PMCID: PMCPMC8288299.

27. Miller HE, Robinson RT. Early control of Mycobacterium tuberculosis infection requires il12rb1 expression by rag1-dependent lineages. Infect Immun. 2012;80(11):3828–41. Epub 2012/08/22. doi: 10.1128/IAI.00426-12. PubMed PMID: 22907814; PubMed Central PMCID: PMCPMC3486065.

28. Case JB, Bailey AL, Kim AS, Chen RE, Diamond MS. Growth, detection, quantification, and inactivation of SARS-CoV-2. Virology. 2020;548:39–48. Epub 2020/08/26. doi: 10.1016/j.virol.2020.05.015. PubMed PMID: 32838945; PubMed Central PMCID: PMCPMC7293183.

29. Robinson RT, Mahfooz N, Rosas Mejia O, Liu Y, Hull NM. SARS-CoV-2 disinfection in aqueous solution by UV222 from a krypton chlorine excilamp. medRxiv. 2021;https://doi.org/10.1101/2021.02.19.21252101 (Preprint posted February 23, 2021).

30. Farkas L, Farkas D, Ask K, Moller A, Gauldie J, Margetts P, et al. VEGF ameliorates pulmonary hypertension through inhibition of endothelial apoptosis in experimental lung fibrosis in rats. J Clin Invest. 2009;119(5):1298–311. Epub 2009/04/22. doi: 10.1172/JCI36136. PubMed PMID: 19381013; PubMed Central PMCID: PMCPMC2673845.

31. Schindelin J, Arganda-Carreras I, Frise E, Kaynig V, Longair M, Pietzsch T, et al. Fiji: an open-source platform for biological-image analysis. Nat Methods. 2012;9(7):676–82. Epub 2012/06/30. doi: 10.1038/nmeth.2019. PubMed PMID: 22743772; PubMed Central PMCID: PMCPMC3855844.

32. Ma D, Chen CB, Jhanji V, Xu C, Yuan XL, Liang JJ, et al. Expression of SARS-CoV-2 receptor ACE2 and TMPRSS2 in human primary conjunctival and pterygium cell lines and in mouse cornea. Eye (Lond). 2020;34(7):1212–9. Epub 2020/05/10. doi: 10.1038/s41433-020-0939-4. PubMed PMID: 32382146; PubMed Central PMCID: PMCPMC7205026.

33. Lafuse WP, Rajaram MVS, Wu Q, Moliva JI, Torrelles JB, Turner J, et al. Identification of an Increased Alveolar Macrophage Subpopulation in Old Mice That Displays Unique Inflammatory Characteristics and Is Permissive to Mycobacterium tuberculosis Infection. J Immunol. 2019;203(8):2252–64. Epub 2019/09/13. doi: 10.4049/jimmunol.1900495. PubMed PMID: 31511357; PubMed Central PMCID: PMCPMC6783358.

34. Jungblut M, Oeltze K, Zehnter I, Hasselmann D, Bosio A. Standardized preparation of single-cell suspensions from mouse lung tissue using the gentleMACS Dissociator. J Vis Exp. 2009;(29). Epub 2009/07/04. doi: 10.3791/1266. PubMed PMID: 19574953; PubMed Central PMCID: PMCPMC2798855.

35. Xin G, Chen Y, Topchyan P, Kasmani MY, Burns R, Volberding PJ, et al. Targeting PIM1-Mediated Metabolism in Myeloid Suppressor Cells to Treat Cancer. Cancer Immunol Res. 2021;9(4):454–69. Epub 2021/02/14. doi: 10.1158/2326-6066.CIR-20-0433. PubMed PMID: 33579728; PubMed Central PMCID: PMCPMC8137571.

